# Community evolution increases plant productivity at low diversity

**DOI:** 10.1101/111617

**Authors:** Sofia J. van Moorsel, Terhi Hahl, Cameron Wagg, Gerlinde B. De Deyn, Dan F.B. Flynn, Debra Zuppinger-Dingley, Bernhard Schmid

**Affiliations:** URPP Global Change and Biodiversity and Department of Evolutionary Biology and Environmental Studies, University of Zürich, Winterthurerstrasse 190, 8057 Zürich, Switzerland; Department of Environmental Sciences, Wageningen University, Droevendaalsesteeg 4,6708 PB Wageningen, the Netherlands

**Keywords:** biodiversity, community evolution, co-selection, ecosystem functioning, grassland species, Jena Experiment, plant productivity, soil organisms

## Abstract

Species extinctions from local communities can negatively affect ecosystem functioning. Ecological mechanisms underlying these impacts are well studied but the role of evolutionary processes is rarely assessed. Using a long-term field experiment, we tested whether natural selection in plant communities increased the effects of biodiversity on productivity. We re-assembled communities with 8-year co-selection history adjacent to communities with identical species composition but no history of co-selection (“naïve communities”). Monocultures and in particular mixtures of two to four co-selected species were more productive than their corresponding naïve communities over four years in soils with or without co-selected microbial communities. At the highest diversity level of eight plant species, no such differences were observed. Our findings suggest that plant community evolution can lead to rapid increases in ecosystem functioning at low diversity but may take longer at high diversity. This effect was not modified by treatments that simulated additional co-evolutionary processes between plants and soil organisms.

## INTRODUCTION

A large number of experiments have shown that species richness positively influences ecosystem functioning, in particular plant biomass production (Tilman *et al.* 1997; Balvanera *et al.* 2006; Cardinale *et al.* 2007, 2012; Reich *et al.* 2012; Meyer *et al.* 2016). These biodiversity effects have been explained by selection effects that increase the chance of including productive species in diverse communities or by complementary effects between species, which allow mixtures to extract resources from the environment more efficiently (Loreau and Hector 2001; Roscher *et al.* 2008; Mueller *et al.* 2013). Furthermore, diversity-dependent reductions in soil fertility (Fornara & Tilman 2008) or density-dependent accumulations of specialist pathogens over time (Schnitzer *et al.* 2011) have been shown to contribute to decreasing productivity at low plant diversity and in plant monocultures.

In contrast to selection effects, complementarity effects between co-occurring species have been shown to increase over time (Cardinale *et al.* 2007; Fargione *et al.* 2007; Reich *et al.* 2012; Meyer *et al.* 2016). Evidence that this might be due to evolutionary processes in plant communities has been found in a glasshouse experiment comparing the community-level performance of plants selected in monocultures vs. plants selected in multi-species mixtures in newly assembled monocultures vs. two-species mixtures (Zuppinger-Dingley *et al.* 2014). This suggests that community evolution may shape diversity-productivity relationship more generally, which could be tested if entire communities of co-selected plant species would be compared with communities of the same plant species but without co-selection history (“naïve communities”). Here, the distinction between the terms “selected” and “co-selected” indicates that in the latter case the outcome of selection on a set of co-occurring species is assessed in the context of a test community formed by these species.

Evolution leading to community-level responses, i.e. changed species performances and interactions due to genetic changes in all or some of the species of the community, has been referred to as community evolution (Goodnight 1990). Later, community evolution was defined as genetically based changes among species constituting the community, which alter species performances and interactions (Wilson 1997; Whitham *et al.* 2006). Such evolutionary changes may occur via genetic recombination, mutations (Anderson *et al.* 2011), or a sorting-out from standing genetic variation through differential survival and growth of individuals (Fakheran *et al.* 2010) and may be due to natural selection or random genetic drift (Wilson & Bossert 1971). In particular, natural selection can lead not only to changes in gene frequencies in populations within species, but selection at the level of communities can in addition lead to correlated changes in gene frequencies in multiple species (Whitham *et al.* 2006) in response to one another or to co-varying environmental conditions. Community evolution can be detected by measurable differences in community-level properties, such as productivity, between communities of co-selected species and naïve communities (Swenson *et al.* 2000; Lawrence *et al.* 2012; Fiegna *et al.* 2014). But empirical evidence for community evolution so far has only been demonstrated in bacterial communities (Lawrence *et al.* 2012; Fiegna *et al.* 2014, 2015) and not yet in higher plants. Here we report results from a field experiment where we tested whether plant community evolution influences plant community productivity.

Recent evidence suggests that selection of particular genotypes from the total genetic pool of a species may affect ecosystem functioning in field experiments (Strauss *et al.* 2008; Lipowsky *et al.* 2011; Lau & Lennon 2012; Kleynhans *et al.* 2016; Rottstock *et al.* 2017). We propose that co-selection of genotypes from the gene pool of entire communities may affect ecosystem functioning if non-random niche or trait changes in response to other phenotypes in the community result in more specialized niches with reduced overlap and a more complete use of biotope space (Dimitrakopoulos & Schmid 2004; Jousset *et al.* 2011), thus leading to increased plant community productivity. We therefore compared the productivity of plant communities assembled from plants that have co-occurred for eight years in a longterm grassland biodiversity experiment (the Jena Experiment, see Roscher *et al.* 2004) with the productivity of plant communities of identical species composition, but without any co-occurrence history (“naïve communities”). The naïve plants were obtained from the seed supplier of the original seeds used to establish the Jena Experiment. We used experimental plant monocultures and 2-, 4- or 8-species mixtures with twelve different species compositions for each diversity level.

To explore if (co-)selection of plant species led to changes in trait distributions within species we used plant height and specific leaf area (SLA) as representative functional traits (Diaz *et al.* 2016), which have been used in previous biodiversity experiments (Roscher *et al.* 2015; Zuppinger-Dingley *et al.* 2014; Cadotte 2017). Variation in these two traits has been shown to affect plant community productivity in contrasting ways: whereas low variation in height is related to asymmetric competition for light and increased productivity via selection effects, high variation in SLA is related to differential light-use strategies and increased productivity via complementarity effects (Roscher *et al.* 2015; Cadotte 2017). As a consequence, we expected a narrowing/widening of the within-species variation in height/SLA at low biodiversity due to community evolution.

Plant community evolution in the field may also depend on the local environment, such as the soils in which co-evolution with soil microorganisms occurs. For instance, plant-soil feedback experiments have shown that soil biota change in response to different plant species, which can in turn modify the composition and productivity of plant communities (Klironomos 2002; Kardol *et al.* 2007). Further, it is thought that negative plant-soil feedbacks may also incur selection for individuals that are able to reduce antagonistic and improve beneficial associations with soil organisms (van der Putten *et al.* 2013; Zuppinger-Dingley *et al.* 2016). To assess whether additional co-evolutionary processes between plants and soil organisms modified plant community evolution, we grew the communities of co-selected plants and naïve communities in soils with co-selected soil organisms (native soil) and with external soil organisms (neutral soil; see Methods and Fig. S1). Community-level plant productivity was measured each year from 2012 to 2015 by collecting species-specific aboveground biomass at the time of peak biomass in spring in each soil treatment, whereas the traits plant height and SLA were measured once in 2015 in the neutral soil only (see Methods).

## METHODS

### Study site

The present study was conducted at the Jena Experiment field site (Jena, Thuringia, Germany, 51°N, 11°E, 135m a.s.l.) from 2011 to 2015. The Jena Experiment is a long-term biodiversity experiment where 60 grassland species have been grown in different combinations since 2002 (Roscher *et al.* 2004).

### Community-evolution treatment (plant history)

The 48 experimental plant communities of this study included twelve monocultures (of which one was removed from all analyses because it was planted with the wrong species), twelve 2-species mixtures, twelve 4-species mixtures and twelve 8-species mixtures. We used two community-evolution treatments (plant histories); plants with eight years of co-selection history in different plant communities in the Jena Experiment (communities of co-selected plants) and plants without such co-selection history (naïve communities). For convenience, we use the term co-selection also for monocultures with a selection history of eight years in the Jena Experiment, even though in this case the term could only apply to potential co-selection with soil organisms. The plant seeds of naïve communities were obtained from the same seed supplier (Rieger Hofmann GmbH, in Blaufelden-Raboldshausen, Germany) as the seeds used for the establishment of the original communities of the Jena Experiment. This supplier collected plants of the different species at field sites in Germany and propagated them for at least five years in monoculture, reseeding them every year. Seeds of communities of co-selected plants were produced in an experimental garden in Zurich, Switzerland, from cuttings that had been made in the Jena Experiment and were then planted in Zurich in the original species combination in plots fenced with plastic netting to reduce pollination between communities. To obtain sufficient numbers of seeds from communities of coselected plants, a small number was additionally collected directly in the plots of the Jena Experiment. All these seeds were thus offspring of plant populations that had been sown in 2002 and grown until 2010 in plots of the Jena Experiment.

The seeds of communities of co-selected plants and naïve communities were germinated in potting soil (BF4, De Baat; Holland) in mid-January 2011 in a glasshouse in Zurich. In March 2011, the seedlings were transported back to the field site of the Jena Experiment and planted within 2 × 2 m subplots of the original plots (Fig. S1). There were four 1 × 1 m quadrats with different soil treatments in each (see next section). Each quadrat was further divided into two 1 × 0.5 m halves. The seedlings of communities of co-selected plants were transplanted into one half and seedlings of naïve communities into the other half of each quadrat at a density of 210 plants per m^2^ (Fig. S1). Species were planted in equal proportions, but if a species was no longer present in an original plot of the Jena Experiment it was excluded from both communities of co-selected plants and naïve communities. Five plant species were excluded in total.

### Soil treatment

Within each 2 × 2 m subplot of the 48 plots of the Jena Experiment used for the present study, the original plant cover was removed in September 2010 and the soil was excavated to a depth of 0.35 m and sieved. To minimize exchange of soil components between quadrats within subplots and with the surrounding soil, two 5-cm layers of sand were added to the bottom of the plots and separated with a 0.5 mm mesh net. The borders of the quadrats and the subplots were separated by plastic frames (Fig. S1). Using the excavated original soil from each of the plots, four soil treatments were prepared. First, half of the soil (approximately 600 kg per plot) was gamma-sterilized to remove the original soil community. Half of the gamma-sterilized soil was then inoculated with 4 % (by weight) of live sugar-beet soil and 4 % of sterilized original soil of the corresponding plot (“neutral soil” obtained by inoculation). Live sugar-beet soil was added to create a natural but neutral soil community and was previously collected in a sugar-beet field not associated with the Jena Experiment, but with comparable soil properties. The other half of the gamma-sterilized soil was inoculated with 4 % (by weight) of live sugar-beet soil and 4 % of live original soil of the corresponding plot (“native soil” obtained by inoculation). The other half of the soil was unsterilized and used for the other two soil treatments. Half of this soil was filled back into one quadrat of the corresponding plot (“native soil”). The other half of the unsterilized soil was mixed among all plots and filled into the remaining quadrats. This fourth soil treatment was abandoned after two years because the plant community was excavated for another experiment. Therefore, this treatment is not included in the present study.

Before the soils were added into the quadrats in December 2010, they were rested in the field in closed bags to allow for the soil chemistry to equalize and to encourage soil biota of the inocula to colonize the sterilized soil before planting. After the soil was added, all quadrats were covered with a net and a water permeable black sheet to avoid spilling between quadrats until the seedlings were transplanted in March 2011.

To test the effectiveness of our soil treatments, we collected soil samples in April 2011 and August 2012 for each quadrat and analysed their fungal communities using terminal-restriction fragment length polymorphism (T-RFLP, see Supporting information). This method allowed us to establish that fungal communities of the soil treatments remained distinct (Table S3). Because such a method does not inform about species identity, the analysis of species-or even clade-specific co-evolutionary interactions between plants and their associated microbial communities was beyond the scope of this study.

### Data collection

We maintained the test communities by weeding three times a year and by cutting the plants twice a year at typical grassland harvest times (late May and August) in central Europe. To measure productivity, we harvested plant material 3 cm aboveground from a 50 × 20 cm area in the centre of each half-quadrat, sorted it into species, dried it at 70°C and weighed the dry biomass.

### Trait measurements

At the end of the experiment, in May 2015, we measured plant height and SLA for 30 species in neutral soil. For each species, we collected up to 20 representative leaves (depending on the leaf size of the species) from four individuals and measured the leaf area by scanning fresh leaves with a Li-3100 Area Meter (Li-cor Inc., Lincoln, Nebraska, USA) immediately after harvest and determining the mass of the same leaves after drying.

### Statistical analysis

We analysed the data from the four spring harvests 2012-2015, which corresponded to peak aboveground plant biomass values. We analysed plant biomass (g/m^2^) as a function of the design variables using mixed models and summarized results in analyses of variance (ANOVA) tables (e.g. Table 1). Significance tests were based on approximate F-tests using appropriate error terms and denominator degrees of freedom. The fixed terms in the model were species richness of the original plots of the Jena Experiment (linear (lSR) and quadratic contrast (qSR) of the logarithm of plant species richness or factor with 4 levels: facSR), year of harvest (linear contrast: linHar), soil treatment (factor with 3 levels: SH), community-evolution treatment (communities of co-selected plants vs. naïve communities: PH as abbreviation for plant history) and interactions of these. The random terms were plot, quadrat, half-quadrat and their interactions with year of harvest. Statistical analyses were conducted using the software R, version 3.2.3 (R Core Team 2015). Mixed models using residual maximum likelihood (REML) were fitted using the package ASReml for R (Butler 2009).

**Table 1.**
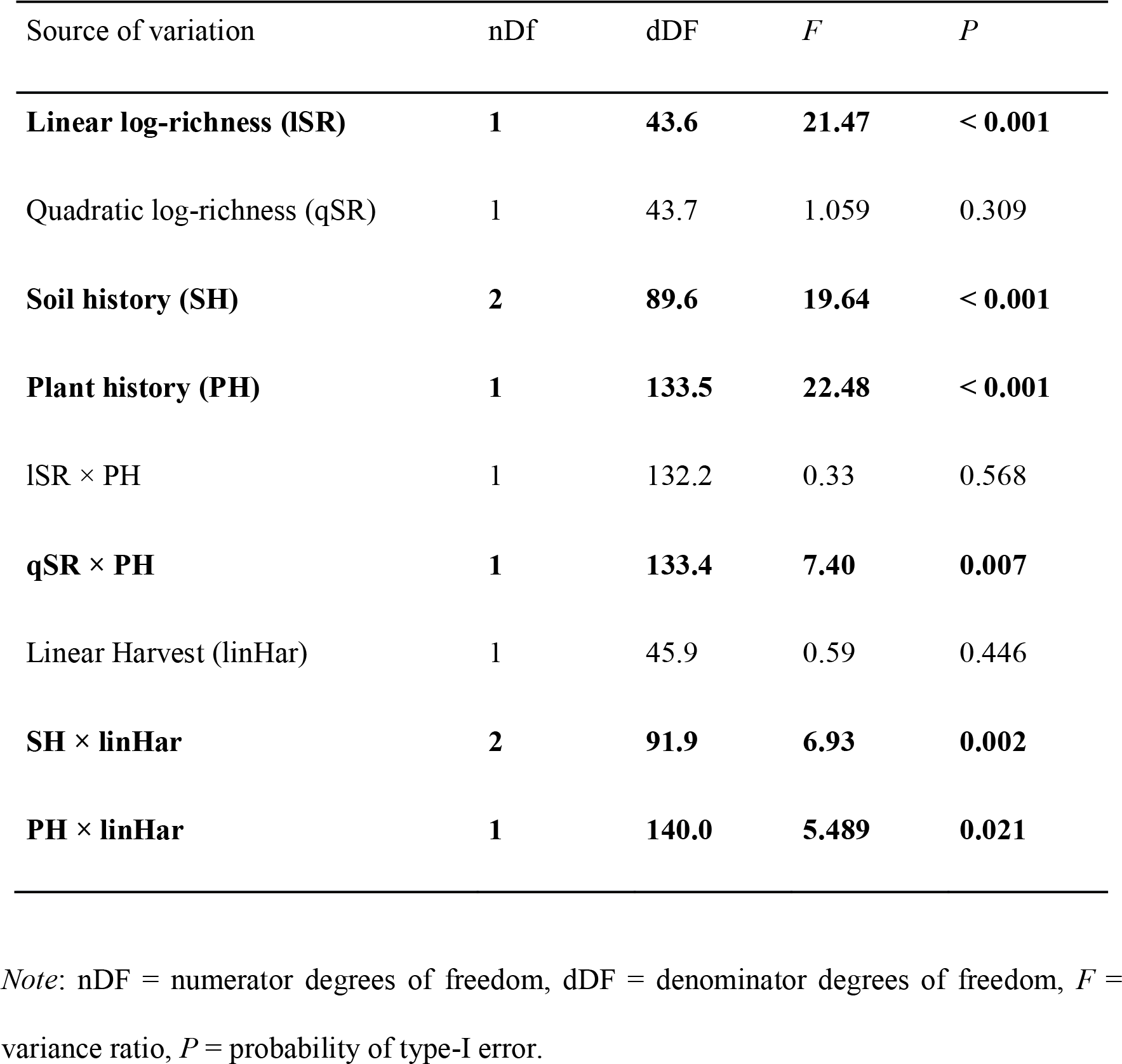
Result of mixed-effects ANOVA for the aboveground biomass of the test communities. Significant effects are shown in bold font.

| Source of variation | nDf | dDF | *F* | *P* |
| --- | --- | --- | --- | --- |
| **Linear log-richness (ISR)** | **1** | **43.6** | **21.47** | **< 0.001** |
| Quadratic log-richness (qSR) | 1 | 43.7 | 1.059 | 0.309 |
| **Soil history (SH)** | **2** | **89.6** | **19.64** | **< 0.001** |
| **Plant history (PH)** | **1** | **133.5** | **22.48** | **< 0.001** |
| ISR × PH | 1 | 132.2 | 0.33 | 0.568 |
| **qSR × PH** | **1** | **133.4** | **7.40** | **0.007** |
| Linear Harvest (linHar) | 1 | 45.9 | 0.59 | 0.446 |
| **SH × linHar** | **2** | **91.9** | **6.93** | **0.002** |
| **PH × linHar** | **1** | **140.0** | **5.489** | **0.021** |

*Note:* nDF = numerator degrees of freedom, dDF = denominator degrees of freedom, *F* = variance ratio, *P* = probability of type-I error.

To compare biomass production between communities of co-selected plants vs. naïve communities across soil treatments and years and to compare these communities from 2012-2015 with the ancestral communities from 2003-2006, we also calculated relative productivities in percentage of the mean productivity of communities at the highest diversity of eight species for each soil treatment x community-evolution treatment x year combination. We chose the 8-species richness level as reference because there were no productivity differences between communities of co-selected plants vs. naïve communities at this diversity level (see Results). Furthermore, we analysed effects of the community-evolution treatment, calculated separately for each particular species composition x soil treatment x year combination, using log-ratios of proportional productivity changes between communities of co-selected plants and naïve communities. In biodiversity experiments, proportional changes typically differ from absolute changes (and absolute changes expressed relative to a control level) because of biomass overyielding at higher compared with lower diversity.

To test whether the presence of particular plant functional groups could differentially affect community evolution, we fitted corresponding contrasts in the analyses of community productivity. We also analysed the responses of individual species to the community-evolution treatment. Differential responses between species could mean that community evolution leads to changes in species abundance distributions, which in turn could lead to changed productivities. We therefore compared the evenness of species aboveground biomasses between communities of co-selected plants vs. naïve communities, species richness levels, soil treatments and years 2012-2015.

Within-species variation in plant height and SLA was calculated as the within-species variance component for each community (residual mean square after fitting species). We had insufficient trait data to test for increased between-species variation in communities of coselected plants containing a mixture of species.

## RESULTS

### Community productivity

Overall, for each doubling of species richness community aboveground biomass increased by 100 g·m^−2^·y^−1^, a typical value for grassland biodiversity experiments (Hector *et al.* 1999) (Fig. 1, Table 1). However, between monocultures and 2- to 4-species mixtures this increase in productivity was steeper for communities of co-selected plants than for naïve communities of the same species composition (*P* < 0.01 for interaction qSR × PH in Table 1). Communities of co-selected plants were also more productive than naïve communities on average. Furthermore, productivity increased across years in communities of co-selected plants but decreased in naïve communities (interaction PH × linHar in Table 1). Because no differences in productivity between the two community-evolution treatments were found at the highest species-richness level (Fig. 1), we also analysed the productivity of all communities in relative terms as percentage of the mean productivity of 8-species mixtures for each plant history x soil treatment x year combination (Fig. 2 and Fig. S2). This analysis confirmed that especially 2- and 4-species mixtures of co-selected plants increased productivity relative to 8-species mixtures (see Supporting Information).

**Figure 1.**
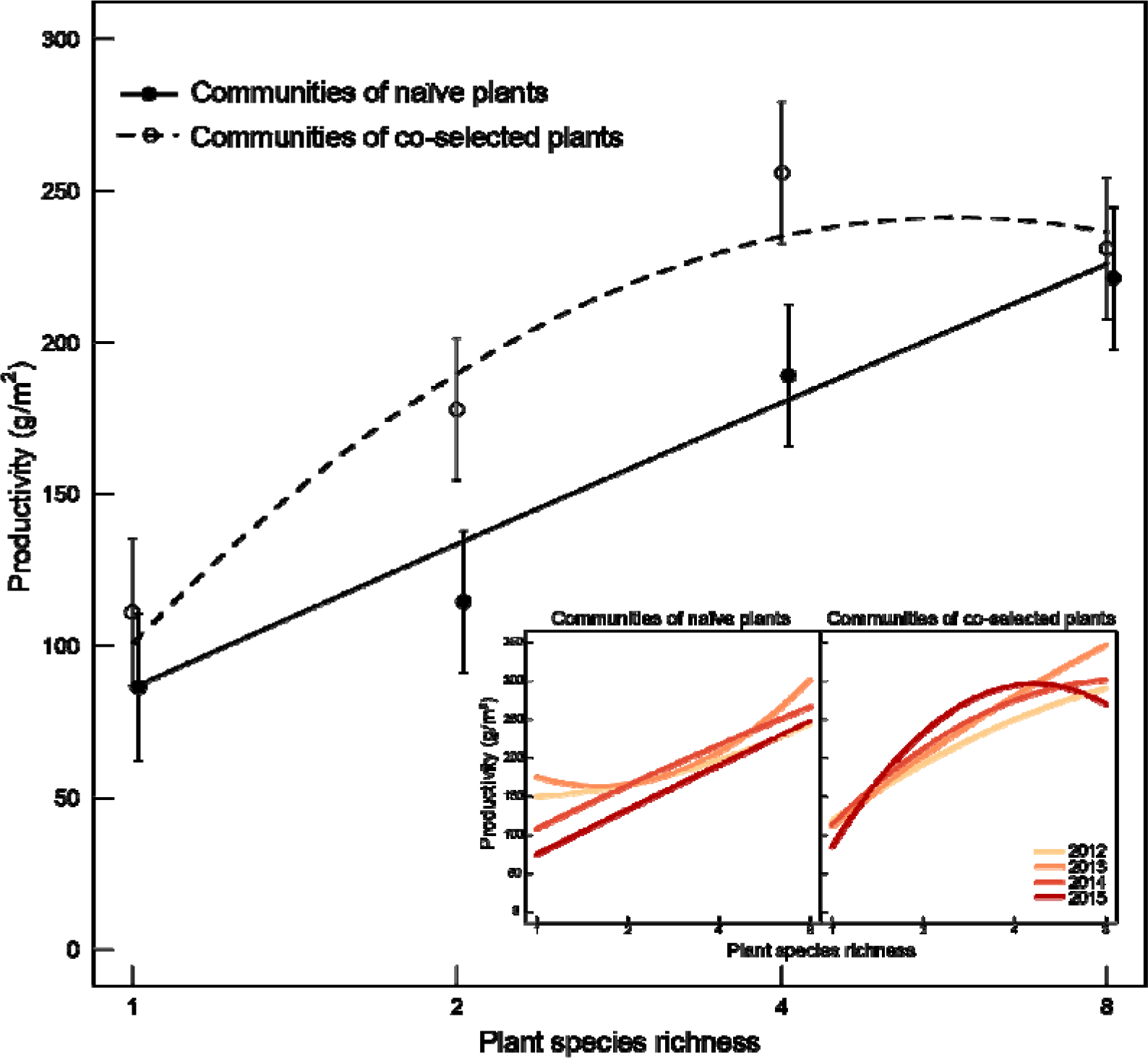
Productivity of communities of co-selected plants (dashed lines, open circles) and naïve communities (solid lines, closed circles). In communities of co-selected plants the increase in productivity from monocultures to 2- and 4-species mixtures was stronger than in naïve plant communities (see Table 1). Main panel: quadratic response curves, averaged over four years. Points are predicted means and standard errors derived from a mixed model including fixed-effects terms for species richness as 4-level factor and community-evolution treatment and random-effects terms for plot, quadrat, half-quadrat and interactions of these with year of harvest (see Methods). Inset panel: separate quadratic response curves for years, with darker colors representing later years.

**Figure 2.**
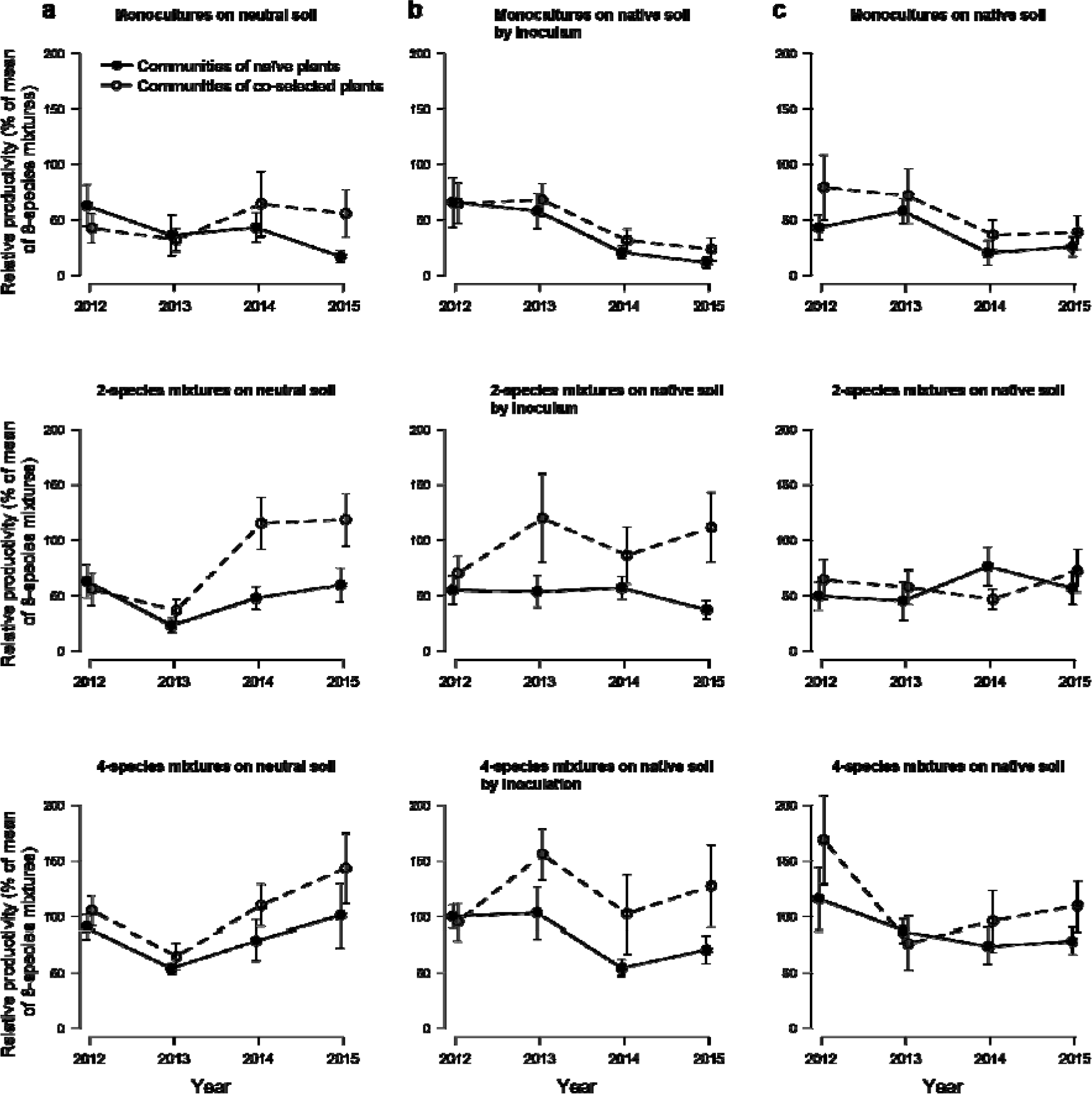
Relative productivity (% of mean of 8-species mixture) of communities of coselected plants (dashed lines, open circles) and naïve communities (solid lines, closed circles) in monocultures and 2- and 4-species mixtures in **(a)** neutral soil (sterilized soil with neutral inoculum) **(b)** native soil obtained by inoculation (sterilized soil with neutral inoculum and inoculum of co-selected soil biota from original plots) and **(c)** native soil (unsterilized soil with co-selected soil biota from original plots). Raw means and standard errors are shown (for significances see Table S1).

### Comparison with ancestor plant communities

To test whether the communities of co-selected plants were already particularly productive in 2- and 4-species mixtures at the beginning of the Jena Experiment (i.e. when they were “naïve” communities themselves), we compared the productivity data of 2003-2006 with the data of 2012-2015. Because productivities were generally higher at the beginning of the Jena Experiment (Fig. S3a), we used relative productivity (percentage of mean of 8-species mixtures per year) to standardize for differences in overall productivity between time periods (Fig. S3b). The plant communities were established on neutral soil in 2002 at the beginning of the Jena Experiment. We therefore used only data from neutral soil for the period 2012-2015. The communities of co-selected plants were significantly different in their response compared to the two types of naïve communities because of their increased relative productivity in 2- and 4-species mixtures (*F*_1,46.5_ = 5.73, *P* = 0.021 for the interaction of plant history with the contrast “2- or 4-species mixtures vs. others”; Fig. S3b). Differences between the communities of the naïve ancestors of the co-selected plants and our current re-assembled naïve plant communities were small and not significant (F_1,46.1_ = 0.23, *P* = 0.637 for the interaction of the contrast “naïve ancestors vs. current naïve communities” with the contrast “2- or 4- species mixtures vs. others”).

### Influence of soil environment

Plant community productivity was initially greater in inoculated soils, which was reflected in an overall main effect of soil treatment and a significant interaction with year (Table 1). This was probably caused by the nutrient flush associated with gamma-sterilization of the soil (Gebremikael *et al.* 2015). But we found no evidence that our soil treatments modified the differences in biodiversity effects between communities of co-selected plants and naïve communities (*P* > 0.2 for the three-way interactions of contrasts of the logarithm of species richness or the 4-level factor of species richness with soil and community-evolution treatments).

### Proportional productivity increases of single species compositions due to community evolution

Consistent with the results of the overall analysis of productivities, the log-ratios of productivities between communities of co-selected plants and naïve communities were on average zero for the 8-species mixtures, that is, the community-evolution treatment did not lead to a proportional increase in productivity at the highest level of species richness (Fig. 3). However, whereas the productivity increases found in the overall analysis were largest in the 4-species mixtures (see Fig. 1), the proportional increases were largest in the 2-species mixtures (Fig. 3). This was due to the overall lower productivity of 2- as compared with 4-species mixtures. Using contrasts between the different diversity levels, we could confirm that the three low diversity levels were significantly different from the 8-species mixtures (*F*_1,37.1_ = 5.34 and *P* = 0.026). Among the three low diversity levels, the 2-species mixtures had significantly greater log ratios than 4-species mixtures and monocultures (*F*_1,39.2_ = 4.44, *P* = 0.042). Comparing 2-species mixtures and monocultures, 2-species mixtures had significantly greater log ratios than monocultures (*F*_1,41.7_ = 4.247, *P* = 0.046).

**Figure 3.**
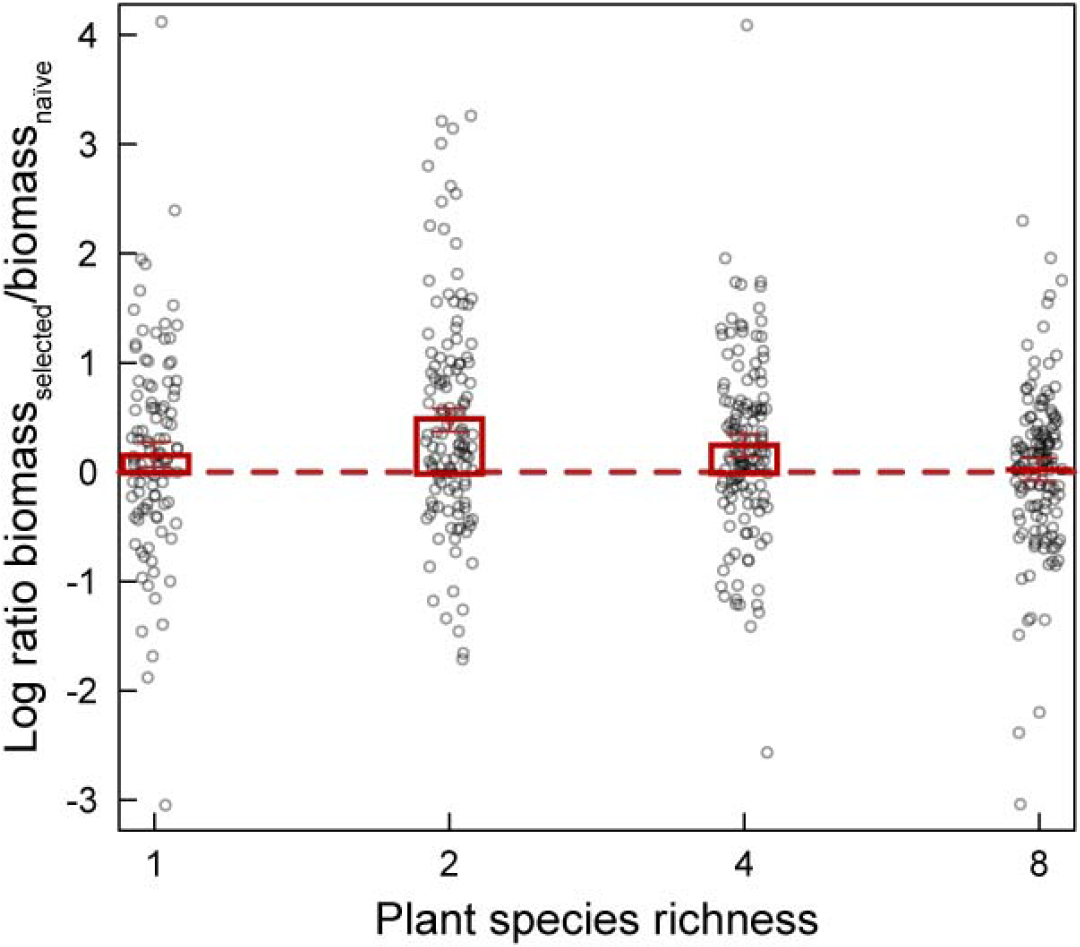
Log ratio of productivity in communities of co-selected plants (bm_selected_) and productivity in naïve communities (bm_naïve_) averaged over four years and three soil treatments. In 8-species mixtures, productivity did not differ between communities of coselected and naïve plants (ratio=0). Especially in 2- and 4-species mixtures, but also in monocultures, communities of co-selected plants produced more biomass than naïve communities. Means and standard errors are shown. Raw data are plotted in the background.

### Influence of functional groups, responses of individual species and evenness

Next, we tested whether the presence of particular plant functional groups influenced the increase in productivity in communities of co-selected plants at the 2- and 4-species richness levels; especially as legumes are known to drive over-yielding in grasslands (Spehn *et al.* 2002). The presence of legumes and other plant functional groups did not provide any further explanation for our results. Species-level productivity within communities was higher for the majority of plant species with a co-selection history, irrespective of functional-group identity (Fig. 4). naïve communities showed marginally more even species abundance distributions (Table S2), mainly due to the lower evenness of communities of co-selected plants in the unsterilized native soil treatment (Fig. S4). Over the course of the experiment, evenness decreased similarly in communities of co-selected plants and naïve communities (Table S2).

**Figure 4.**
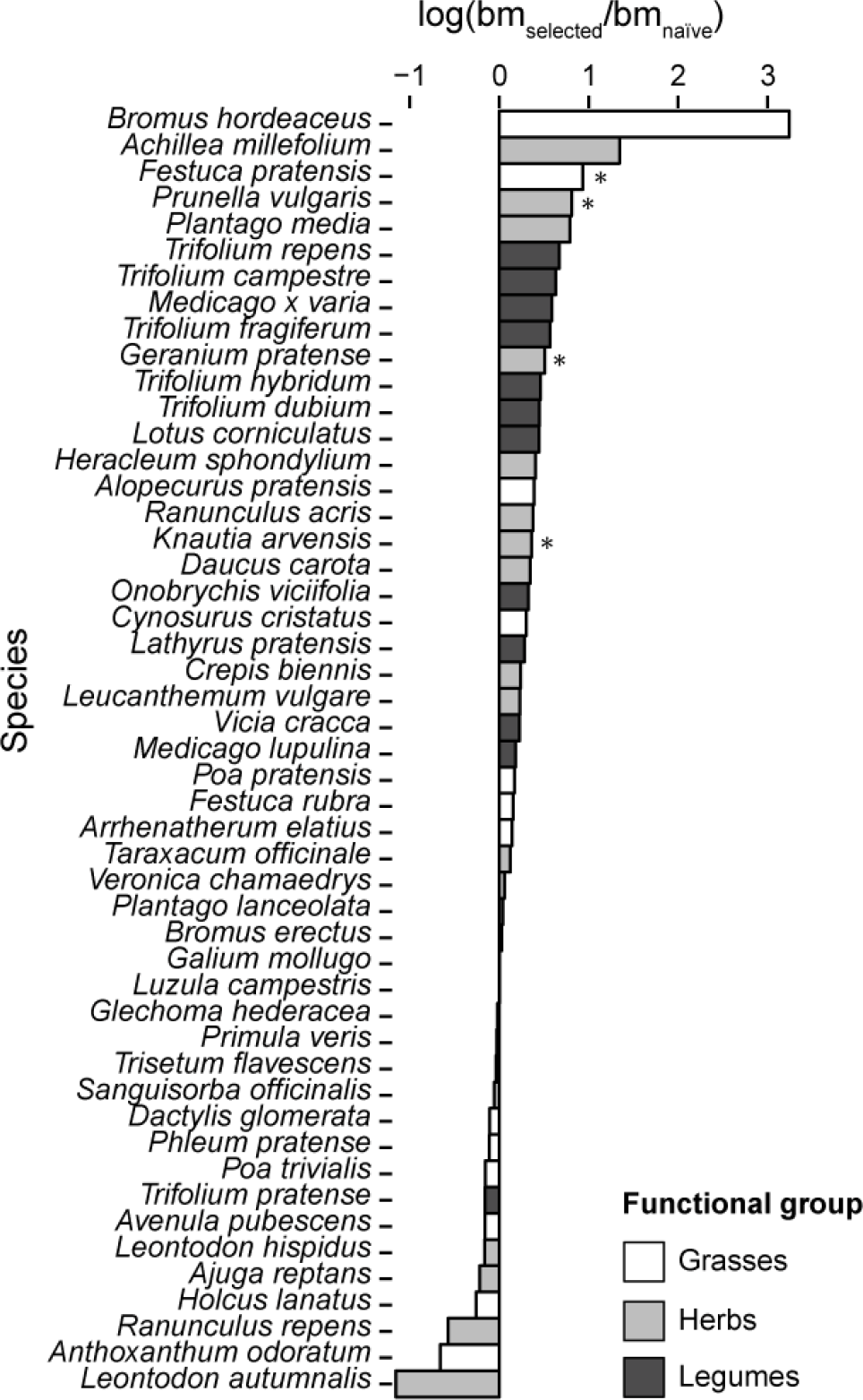
Log-transformed species biomass ratios between co-selected and naïve plants. The majority of plant species attained greater aboveground biomass in communities of co-selected plants compared with naïve communities. The studied plant species belong to three different functional groups: grasses (white bars), herbs (light grey bars) and legumes (dark grey bars). Data are for each species averaged over four years, three soil treatments, four species-richness levels and species compositions within richness levels (*n* = 32-352). Three species with n < 32 were excluded from the analysis (*Anthriscus sylvestris, Campanula patula* and *Cardamine pratensis*). The stars represent *P*-values < 0.05 for species tested separately.

### Within-species trait variation

Finally, we analysed changes in within-species trait variation along the species richness gradient as a potential mechanism contributing to the difference in productivity between communities of co-selected plants and naïve communities (Roscher *et al.* 2015; Cadotte 2017). Within-species variation in plant height was marginally reduced at low diversity in communities of co-selected plants compared with naïve communities (Fig. 5a; *F*_1,733_ = 3.187, *P* = 0.078 for the interaction of log species richness with plant history). In contrast, within-species variation in specific leaf area (SLA) was increased at low diversity for communities of co-selected plants and decreased for naïve communities (Fig. 5b; *F*_1,69.2_ = 4.87, *P* = 0.031 for the interaction of log species richness with plant history).

**Figure 5.**
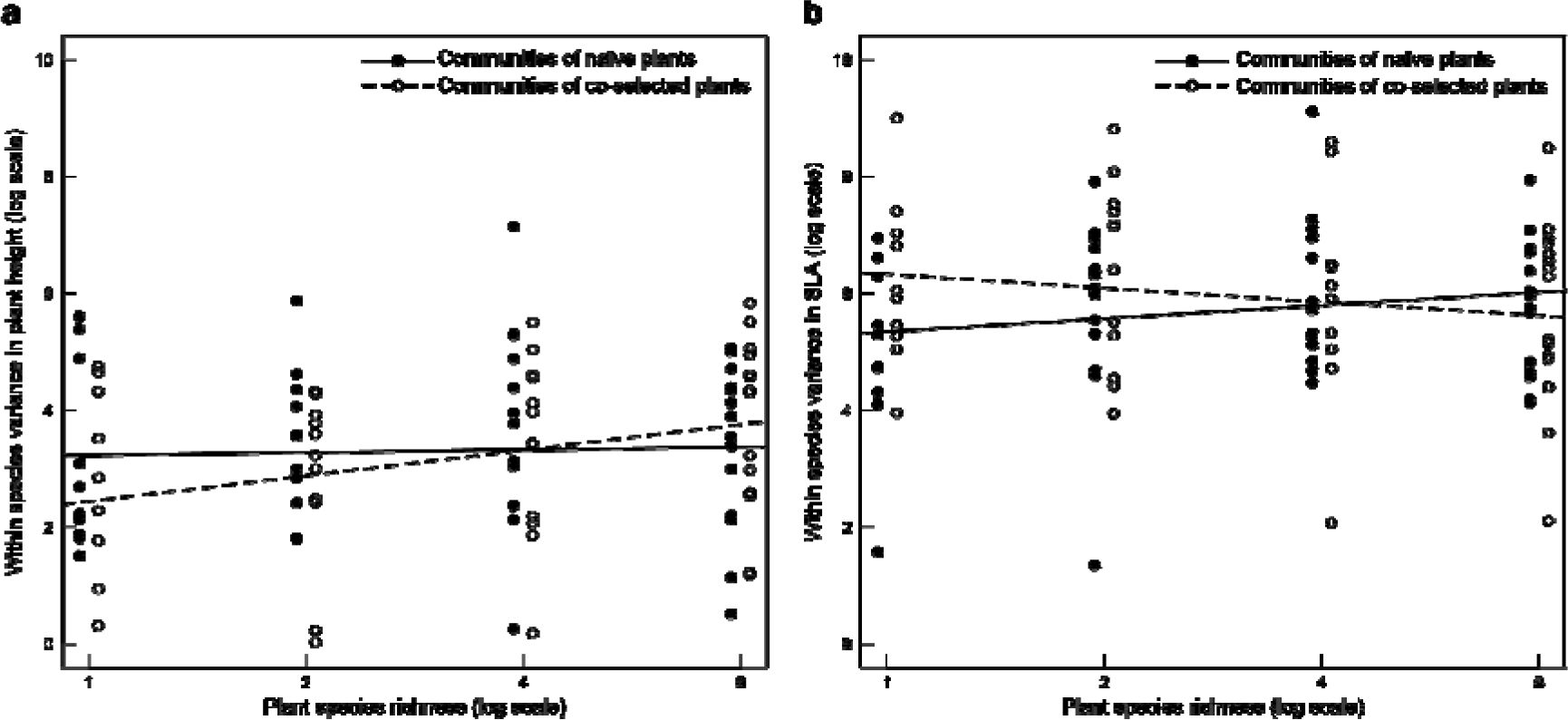
Within-species variation in plant height **(a)** and specific leaf area (SLA) **(b)** for communities of co-selected plants and naïve communities at the end of the experiment in 2015 in neutral soil. In monocultures within-species variations in height/SLA (measured as the within-species variance component in analysis of variance) were smaller/larger for coselected than for naïve plants and these differences decreased with increasing species richness. Open circles and dashed line refer to communities of co-selected plants, closed circles and solid line refer to naïve communities.

## DISCUSSION

Our results show that eight years of community evolution in a biodiversity experiment can increase biodiversity effects on community productivity, suggesting that this may at least in part explain why biodiversity effects commonly increase over time in such experiments (Cardinale *et al.* 2007; Fargione *et al.* 2007; Reich *et al.* 2012; Meyer *et al.* 2016). The greater productivity in communities consisting of co-selected plants compared with communities consisting of naïve plants was particularly evident in communities comprised of two or four species whereas 8-species mixtures with and without co-selection history on average showed the same productivity. It is conceivable that selection pressure was dampened in communities where more than four species co-occurred. For instance, during initial establishment in a diverse community, each individual can have a very different set of immediate neighbours that could constrain the consistency in the selection pressure on individuals within a community. With fewer species in a mixture, the potential for the evolution of increased complementarity between plant species should be greater, given the relative constancy of the neighbours any given plant experiences. The greater proportional increase of productivity in communities of co-selected plant species at the 2- than at the 4-species richness level, and the absence of such an increase at the 8-species richness level, are compatible with the idea that evolution for co-adaptation is stronger at low than at high diversity. As suggested by Cardinale et al. (2012), there might be an upper limit of species richness beyond which selection is unlikely to strengthen biodiversity effects. Additionally, community evolution leading to increased plant growth and productivity in diverse mixtures may be at the expense of reduced pathogen defence (Lemmermeyer *et al.* 2015).

The performance of the naïve communities in the current study over the four years was comparable to the initial performance of the ancestral community of the co-selected plants (2003-2006). This similarity supports the view that the observed results at 2- and 4-species richness levels in communities of co-selected compared with communities of naïve plants are likely due to diversity-dependent community evolution. Indeed, the naïve communities did not catch up with the communities of co-selected plants during the course of the current experiment and differences in productivity from 2012 to 2015 even increased between the two community-evolution treatments (see Fig. 2). This suggests that in our study community evolution was not solely due to an immediate sorting out of genotypes from standing variation within species (Fakheran *et al.* 2010) during seedling establishment and initial growth. However, differential responses of individual species to the community-evolution treatments (see Fig. 4) could have caused the marginally decreased evenness of species abundance distributions in communities of co-selected plants. Thus, abundance shifts between species could have contributed to the higher productivities of communities of co-selected plants at intermediate diversity levels or they might have contributed to avoid reduced productivities of naïve communities at the highest diversity level. A further driving force behind community evolution for greater productivity at low diversity may have been strong responses of particular functional groups or functional group combinations to co-selection (Zuppinger-Dingley *et aL.* 2014). There was, however, no evidence for any functional-group specific effect typically found in other contexts of biodiversity-ecosystem functioning research (Hooper & Vitousek 1997; Spehn *et al.* 2002).

Intraspecific variation in plant height was smaller and intraspecific variation in SLA was larger at low diversity in communities of co-selected plants than in naïve communities (see Fig. 5), a result in line with previous findings regarding effects of interspecific variation in these traits on community productivity (Roscher *et al.* 2015; Cadotte 2017): because plant-height related competition for light is asymmetric variation in this trait does not increase complementarity and productivity in plant stands but variation in SLA can do so because it is related to differential light-use strategies. Our results suggest an evolutionary narrowing of height-related niches and an evolutionary broadening of SLA-related niches in monocultures and low-diversity mixtures in (co-)selected plants. At higher diversity this trend was reversed (see Fig. 5), perhaps because here intraspecific variation is relatively less important than interspecific variation. For example, the narrowing of within-species variation in SLA with increasing diversity in communities of co-selected plants is an expected consequence of character displacement between species (Zuppinger-Dingley *et al.* 2014). In contrast, the increasing within-species variation in SLA with increasing diversity in naïve communities (see Fig. 5b) may have been caused by a more heterogeneous biotic environment at high diversity. Because our assessment of plant functional traits was limited to two traits measured at the end of the experiment, the above interpretations can only hint at some mechanisms potentially underpinning the observed effects of our community-evolution treatment on community productivity. As pointed out for example by Cadotte (2017), multivariate trait variation may be more important than variation in any particular trait in particular for complementarity effects.

Our community-evolution treatment also led to increased productivity in monocultures, where plants could not have been co-selected with other plant species but only with soil organisms. This would be consistent with an earlier finding from the Jena Experiment that showed a reduction of negative plant-soil feedbacks in plants that had been grown for 8 years in monoculture compared with plants that had been grown in mixtures (Zuppinger-Dingley *et al.* 2016). There was a slight indication that naïve plants where initially less negatively affected in neutral than in native soil but over time showed increased negative effects (see Fig. 2), possibly due to accumulation of soil pathogens (Schnitzer *et al.* 2011). An alternative but not mutually exclusive explanation for the higher productivity of monocultures of (co-)selected plants than naïve plants could be the evolution of increased resource-uptake niches within species as indicated in the previous paragraph (Bazzaz 1996): assuming a correlation between resource-uptake and trait-based niches (Roscher *et al.* 2015), the increase in within-species variation in SLA (and the decrease in within-species variation in plant height) in monocultures of selected plants would be consistent with this explanation.

Positive plant diversity-productivity relationships may not only be driven by complementary resource use, and thus increased performance at high diversity (Roscher *et al.* 2008; Mueller *et al.* 2013), but also by pathogen accumulation in the soil and thus reduced performance at low diversity (Schnitzer *et al.* 2011). Previous studies in the context of biodiversity-ecosystem functioning research have reported negative plant-soil feedbacks in native as opposed to neutral soils (Klironomos 2002; Petermann *et al.* 2008; Cortois *et al.* 2016). Consequently, an increase of biodiversity effects during community evolution could also be due to the presence of co-selected soil biota. In our study, however, the outcome of the community-evolution treatment in mixtures was largely independent of the presence of co-selected soil biota. It is conceivable that co-evolution of plants with soil biota in our experimental systems was not effective because the large population sizes and short generation times of most soil organisms contributed to the re-assembly and fast evolution of soil communities (Lau & Lennon 2012).

Changes in the performance of individual species selected at different species-richness levels and tested under experimental abiotic or biotic conditions have been observed in previous studies (Lipowsky *et al.* 2011; Zuppinger-Dingley *et al.* 2014; Kleynhans *et al.* 2016; Rottstock *et al.* 2017). Here, we demonstrated that changes in the performance of entire plant communities over time depend on a history of co-selection among the plant species of the assembled mixtures. We suggest that these changes are the result of community evolution because they were maintained through seed production in an experimental garden and propagation of seedlings in a glasshouse to the replanting of communities in the field. However, we cannot exclude maternal carry-over and epigenetic changes (Verhoeven *et al.* 2016) as additional potential evolutionary mechanisms. The observed effects of our community-evolution treatment could have been due to co-selection between the species within each particular community composition. Alternatively, in a more general process, “diffuse” co-selection may have increased the ability of species to grow with any other species. Considering the large residual variation among the productivity responses of the particular community compositions (see Fig. 3), a mixture of mechanisms seems likely. Nevertheless, that the naïve plants were generally poorly adapted to grow with other species or under the abiotic conditions at the field site would not be consistent with their equally high community productivity as that of co-selected plants in 8-species mixtures.

Independent of the mechanism, our findings of positive effects of community evolution on community productivity suggest that for the conservation of biodiversity and its beneficial influence on ecosystem functioning and services it may be best to preserve plant species in a community context rather than in separation from other plant species (and soil biota). Where only the latter is possible, we recommend that populations from communities of particular species compositions are nevertheless kept separate to later re-assemble these communities with those populations that have evolved together.

## ACKNOWLEDGEMENTS

Thanks to the Jena Experiment for providing infrastructure and help, to D. Trujillo and M. Furler for technical assistance and to H. Martens for the lab work for the T-RFLP analyses. Thanks to V. Yadav for the establishment of the plots. We thank Tim Paine, Marc Cadotte and Mark van Kleunen for their helpful comments on the manuscript and Akira Mori and an anonymous reviewer for their very constructive comments during the reviewing process. This study was supported by the Swiss National Science Foundation (grant numbers 130720, 147092 and 166457 to BS), the Netherlands Organisation for Scientific Research (NWO-ALW Vidi grant to GBDD) and the University Research Priority Program Global Change and Biodiversity of the University of Zurich. The Jena Experiment is funded by The Deutsche Forschungsgemeinschaft (DFG, German Research Foundation, FOR1451).

## AUTHORSHIP

BS, DFBF and GBDD conceived the project; DZ-D set up the experiment; SJVM, TH and DZ-D carried out the experiment; BS, CW, SJVM and TH analysed the data; DFBF analysed the TRFLP data; BS, SJVM, TH and CW wrote the first draft of the manuscript. All authors contributed to the final manuscript.

## SUPPORTING INFORMATION

Additional Supporting Information may be found online in the supporting information tab for his article. References unique to the Supporting Information appear only there.

